# Malaria parasites differentially sense environmental elasticity during transmission

**DOI:** 10.1101/2020.09.29.319020

**Authors:** Johanna Ripp, Jessica Kehrer, Xanthoula Smyrnakou, Nathalie Tisch, Carmen Ruiz de Almodovar, Friedrich Frischknecht

## Abstract

Transmission of malaria-causing parasites to and by the mosquito rely on active parasite migration and constitute bottlenecks in the *Plasmodium* life cycle. Parasite adaption to the biochemically and physically different environments must hence be a key evolutionary driver for transmission efficiency. To probe how subtle but physiologically relevant changes in environmental elasticity impact parasite migration, we introduce 2D and 3D polyacrylamide gels to study ookinetes, the parasite forms emigrating from the mosquito blood meal and sporozoites, the forms transmitted to the vertebrate host. We show that ookinetes adapt their migratory path but not their speed to environmental elasticity and are motile for over 24 hours on soft substrates. In contrast, sporozoites evolved more short-lived rapid gliding motility for rapidly crossing the skin. Strikingly, sporozoites are highly sensitive to substrate elasticity possibly to avoid adhesion on soft endothelial cells on their long way to the liver. Hence the two migratory stages of *Plasmodium* evolved different strategies to overcome the physical challenges posed by the respective environments and barriers they encounter.

**Highlights:** *Plasmodium* ookinetes can move for over 24 hours on very soft substrates mimicking the blood meal

*Plasmodium* ookinetes change their migration path according to substrate stiffness

*Plasmodium* sporozoites are highly sensitive to subtle changes in substrate elasticity

Sporozoite may have evolved to not attach to the soft endothelium to help reach the liver

## Introduction

*Plasmodium* parasites need to invade host cells and migrate across tissues at different stages throughout their life cycle (**Figure 1 A**). Their substrate-dependent form of locomotion, termed gliding motility, allows for very fast cell migration that does not involve any cellular protrusions and is propelled by an actomyosin motor (1). The parasite forms developing inside the mosquito midgut after an infectious blood meal, the ookinetes, have to leave the blood meal and traverse the midgut epithelium to transform into oocysts at the basal membrane (2–5). These ookinetes already move actively within the soft blood meal before migrating into the epithelium (6, 7). Inside the oocyst, sporozoites are formed which can infect vertebrates during a second blood meal of the mosquito. When fully developed, these sporozoites move inside the oocyst and egress (8). They are transported within the mosquito circulatory system until they attach to and invade the salivary gland (9). There is only little movement inside the gland but once the sporozoites are transmitted into the skin, they start to migrate at high speed within the dermis (10–12). When a sporozoite finds a blood capillary, it can enter the blood stream, which transports the sporozoites to the liver (13, 14). After exiting the circulation and traversal of several hepatocytes, the sporozoite finally invades a hepatocyte and develop into thousands of red blood cell infecting merozoites (14, 15). How the motile *Plasmodium* stages recognize their target tissues and how the environment influences motility is largely unknown. Several sporozoite proteins have been found to be important for salivary gland invasion (16). Whether there is a specific receptor mediating tissue-specific invasion of salivary glands is currently under debate (17, 18). Liver invasion is thought to be triggered by highly sulphated heparan sulphate proteoglycans which protrude through fenestrations of endothelial cells lining the liver sinusoids (14, 19, 20).

**Figure 1.**
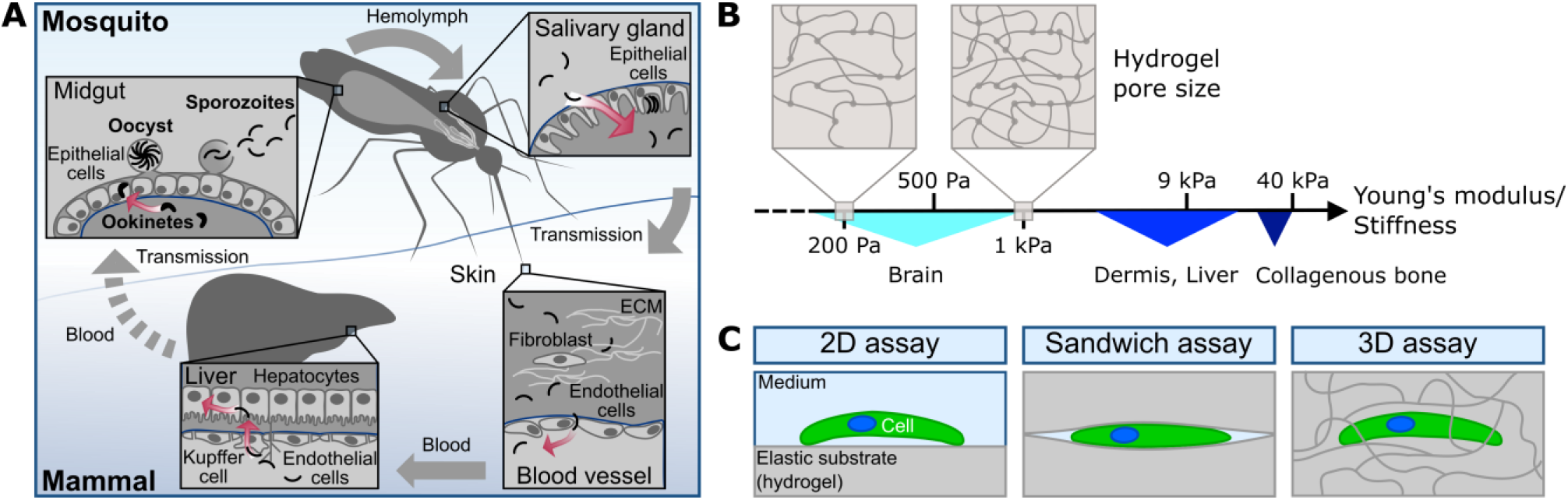
*Plasmodium* parasites need to traverse and invade different host cells and tissues to progress through the life cycle. (A) Ookinetes develop in the midgut and traverse the midgut epithelium to transform into an oocyst at the basement membrane. After rupture of the oocyst, sporozoites are released into the hemolymph. They invade the mosquito salivary glands and move fast upon injection into the dermis via the mosquito bite. Sporozoites enter blood vessels, are transported with the blood circulation, attach to endothelial cells and enter the liver tissue until they finally invade a hepatocyte for transformation and proliferation. Barriers faced by the parasites are depicted in blue. Red arrows indicate active migration. (B) Polyacrylamide hydrogels can be fabricated to mimic the stiffness of different tissues by adjusting the monomer to crosslinker concentration. A higher amount of crosslinker leads to smaller pores and higher elastic moduli of the gels. (C) Experimental setups used in this study to investigate the effect of substrate elasticity, confinement and pores size on ookinete and sporozoite motility.

*In vivo* imaging has been used to characterize *Plasmodium* motility in its natural environment, but it does not allow the tuning of individual biochemical or mechanical properties. In contrast, various *in vitro* assays allowed to examine the influence of defined environmental factors such as ligand-density, hydrodynamic flow, substrate topography and elasticity on sporozoite motility (21–24). This revealed that sporozoites move best on stiff substrates with intermediate ligand density (23), follow environmental topographic cues on their way through the skin (22) and slow motility under flow (21). However, these studies were limited to 2D surfaces, were not suited to investigate ookinetes and failed to compare the small but possibly important elasticity differences relevant for cells encountered by sporozoites. Indeed, the elasticity of human tissues varies from soft brain tissue to intermediate dermal to stiff bone tissue (25, 26) with endothelial cells lining the interior of blood vessels being softer than dermal fibroblasts (27) (**Figure 1 B**).

Here, we introduce tunable polyacrylamide (PA) hydrogels to investigate *Plasmodium* ookinete and sporozoite motility on and within elastic 2D and 3D substrates (**Figure 1 C**). As opposed to natural hydrogels such as matrigel or collagen, PA hydrogels can be manufactured with defined elasticity and pore size by adjusting the monomer to crosslinker concentration (28–30). This allowed us to mimic different microenvironments and study the effect of single physiologically relevant physical parameters on motility of wild type and mutant parasites. We show that ookinetes change their migration path to disseminate faster on stiffer substrates, can actively migrate for over one day while short lived sporozoite motility in 3D hydrogels mimics their migration in the skin. Importantly, we find that the elastic nature of endothelial cells limits the capacity of sporozoites to move on them, suggesting that sporozoites evolved to avoid adhesion to soft endothelial cells in order to ensure long-distance dispersion within the blood flow and efficient homing to the liver.

## RESULTS

### Substrate stiffness impacts ookinete migration pattern

To investigate ookinete motility we generated non-adhesive PA hydrogels with various elasticities (25, 28). We indeed observed reduced adhesion of ookinetes to these non-coated planar PA hydrogels and no continuous movement. We therefore sandwiched the parasites between two hydrogels to establish a confined environment (**Figure 2 A**). This confinement was sufficient to induce ookinete motility in the absence of specific ligands (**Supplementary Movie 1**). As opposed to experiments on glass, where only about half of the ookinetes moved, nearly all ookinetes were motile if sandwiched between two elastic PA hydrogels independent of gel elasticity (**Figure 2 B**). The speed was enhanced 2-3–fold on elastic PA hydrogels compared to glass with only minor differences between soft and stiff hydrogels (**Figure 2 C**). Interestingly, ookinetes were moving for over 20 hours on hydrogels, while they already stopped moving on glass within about three hours (**Figure 2 D**). After 20-26 hours of motility, ookinetes moved more robustly on soft than on stiff gels but were fastest on stiff gels (**Figure 2 D, E**). Strikingly, ookinetes were mostly moving in circular trajectories on soft hydrogels while they were moving more directional on stiffer substrates (**Figure 2 F**). This resulted in a higher mean square displacement of ookinetes moving on stiffer hydrogels (**Figure 2 G**). We hypothesize that this switch in behavior might play a role *in vivo* as ookinetes move out of the soft blood bolus and encounter a stiffer layer of epithelial cells or once the ookinete has passed through the cells and faces the basal lamina (3, 31).

**Figure 2.**
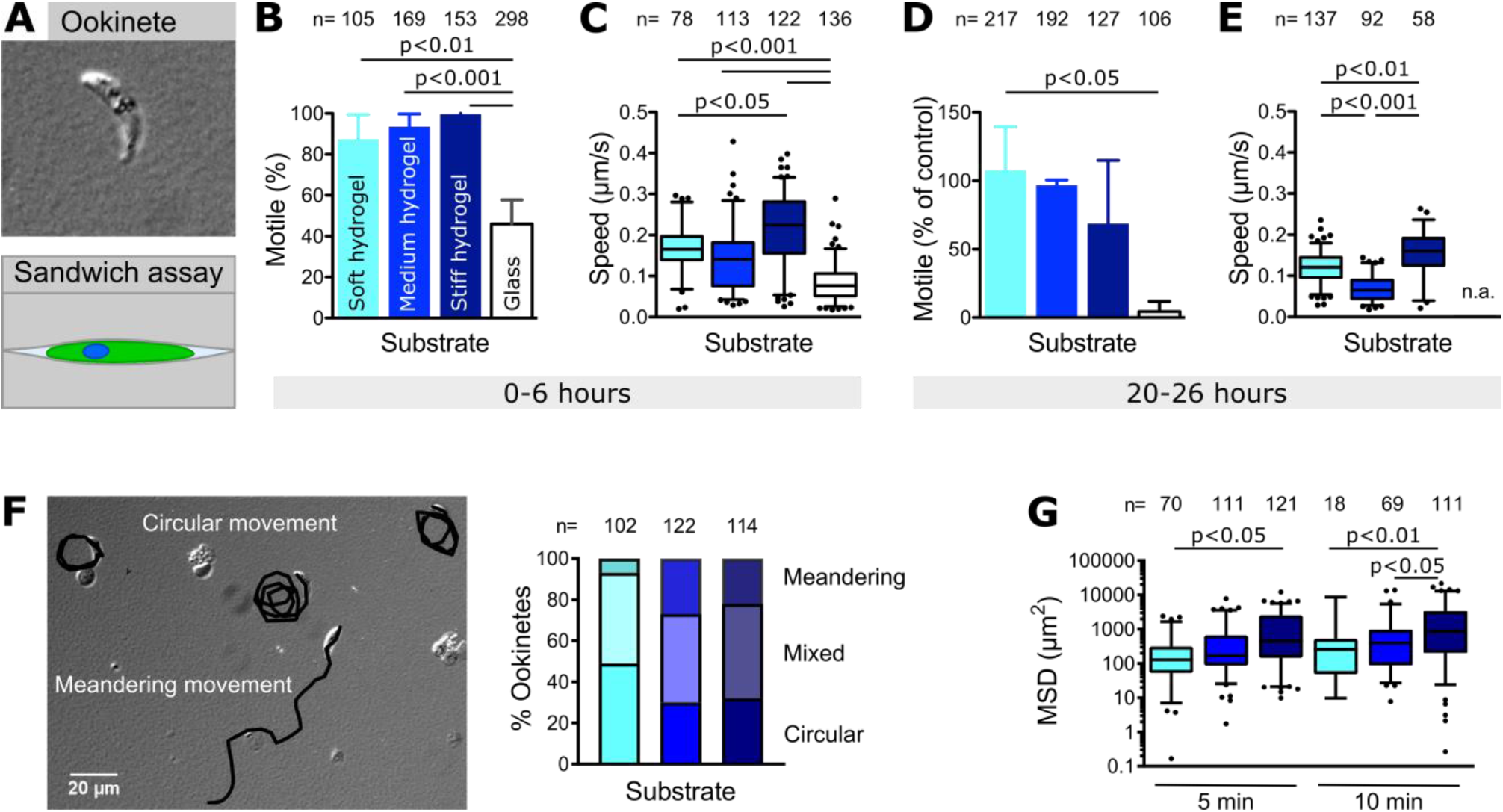
Substrate stiffness affects motility patterns of ookinetes. (A) Ookinetes sandwiched between two hydrogel surfaces. (B) Percentage of ookinetes moving for more than one parasite length within the time of observation (5-10 min) in different sandwiches as indicated. Bars represent the mean and error bars the standard deviation of at least three independent experiments. (C) Speed of motile ookinetes on substrates of different stiffnesses as indicated in panel B. Box-and-whisker plots depict the 25% quantile, median, 75% quantile and nearest observations within 1.5 times the interquartile range (whiskers). Outliers beyond this range are shown as black dots. (D) Percentage of motile ookinetes after 20-26 hours of incubation between hydrogels compared to parasites directly imaged after setting up the experiment as shown in (B). (E) Speed of motile ookinetes after 20-26 hours of incubation between hydrogels. Speed on glass was not analyzed (n.a.) due to the low fraction of motile ookinetes at this timepoint. (F) Migration patterns of ookinetes. Image: Overlay of tracked migration paths (black) and DIC image showing ookinetes at the end of the recorded image sequence. Graph: Quantitative analysis of the indicated different migration patterns on soft, medium and still hydrogels. (G) Mean square displacement of ookinetes moving on hydrogels of different stiffnesses at two timepoints. Numbers above indicate the number of ookinetes analyzed. Significance for (B) and (D) determined by One-way analysis of variance with Bonferroni’s Multiple Comparison test. Significance for (C), (E) and (G) determined by Kruskal-Wallis test with Dunn’s Multiple Comparison test.

### Reconstructing skin-like sporozoite migration in tunable 3D hydrogels

Next, we probed if PA hydrogels could be used for the study of motile sporozoites in 3D. Large metazoan cells do not enter into synthetic hydrogels as they cannot degrade these gels or squeeze through the pores. In line with this, we did not observe any ookinetes penetrating the PA hydrogels. However, when we sandwiched infected salivary glands between a soft PA hydrogel and a glass coverslip, we observed sporozoites moving into the hydrogel (**Figure 3 A**). Inside these hydrogels they mainly moved in helical trajectories (**Figure 3 B, Supplementary Movie 2**) similarly to what has been observed in matrigel (32). While some sporozoites moved horizontally to the plane of image acquisition, others moved vertically and could not be tracked over time as they were quickly moving in and out of focus; some also stopped moving. To study the effect of pore size on sporozoite motility, we manufactured soft hydrogels with different concentrations of crosslinker, thereby tuning the pore size (28, 30). With decreasing pore size, sporozoites slowed down and a higher fraction of sporozoites seemed to get stuck in the hydrogel and eventually sporozoites failed to enter the hydrogels. The motility activating bovine serum albumin had to be added to the medium for sporozoites to move inside the hydrogels, but not to enter the hydrogels. Importantly, sporozoite speed in both hydrogels was similar to the speed distribution observed in the skin *in vivo*, with the sporozoites in the small pore gel representing the slower *in vivo* fraction, while the sporozoites in the large pore gel were representing the faster fraction (**Figure 3 C**). To test, how accurately soft PA hydrogels can be used to mimic sporozoite migration in the skin we analyzed two transgenic parasite lines respectively lacking the actin-binding protein coronin (33) and the heat shock protein 20 (HSP20) (34). We chose these lines as they show different motility phenotypes in the skin and on glass: *coronin(-)* sporozoites moved well in the skin but showed a migration defect on glass (33). In contrast, *hsp20(-)* sporozoites moved slower both on glass and within the skin (34). Interestingly, *coronin(-)* sporozoites moved in PA hydrogels with small pores as well as wild type sporozoites and almost as well in the PA hydrogel with large pores (**Figure 3 D, E**). In contrast, *hsp20(-)* sporozoites moved inefficiently in both gels (**Figure 3 D, E**). This shows that sporozoite migration parameters in the PA hydrogels largely mimicked those observed in the skin in all three investigated parasite lines. To further test the versatility of the gel system, we assessed if drugs can diffuse into the hydrogel and affect motility. Indeed, the actin polymerization inhibitor cytochalasin D (CytoD), which inhibits migration *in vitro* (35), also reduced the speed of sporozoites moving in PA hydrogels at a low concentration of 25 nM (**Figure 3 F, G**), while at 50 nM sporozoites did not enter the gels. Together, these data suggest that the 3D PA hydrogels can be used to mimic key features of sporozoite migration in the skin and can serve as model system to rapidly test drugs or antibodies against the parasite during the first minutes of a *Plasmodium* infection.

**Figure 3.**
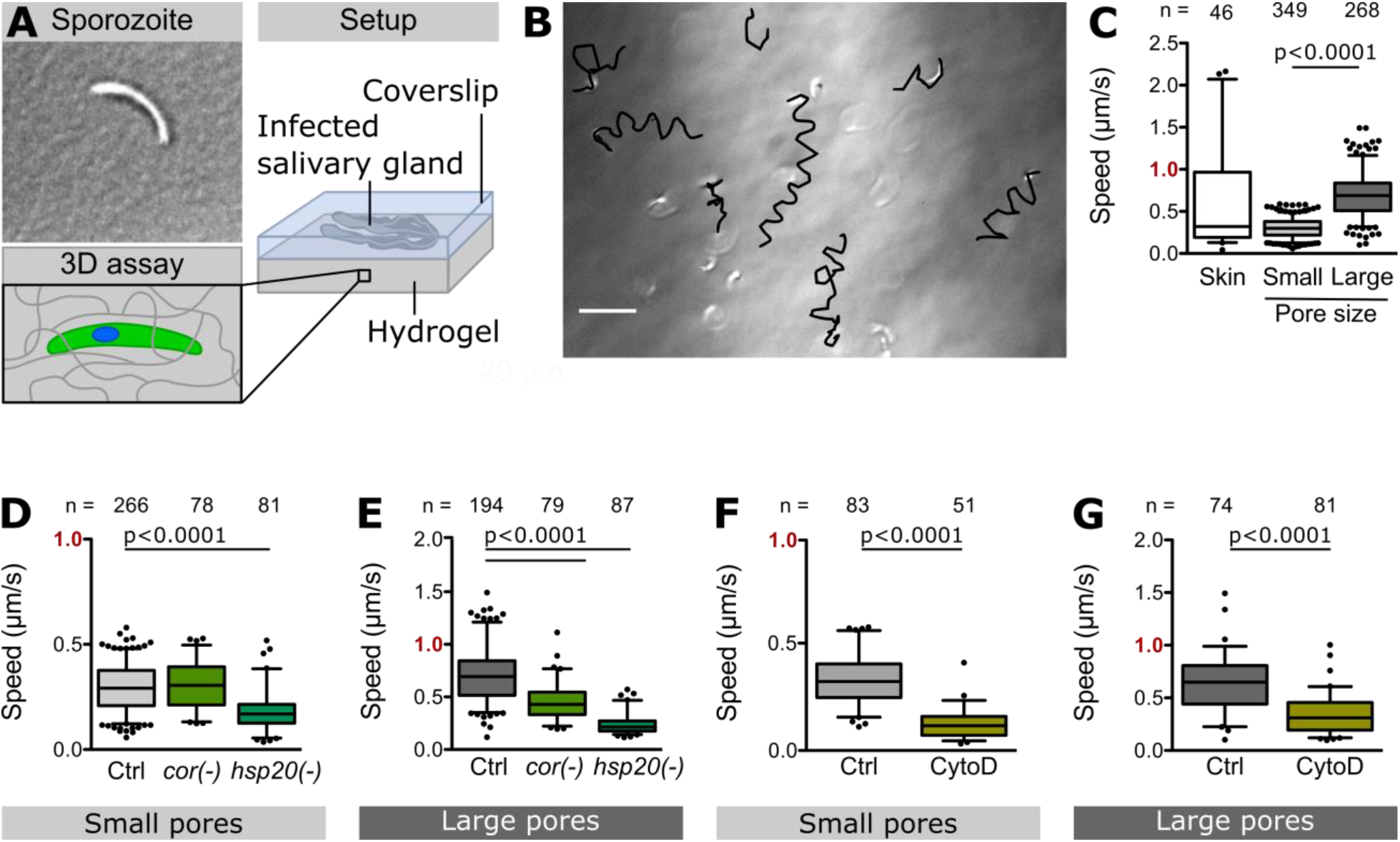
Sporozoite motility inside 3D hydrogels. (A) Infected salivary glands were sandwiched between a soft PA hydrogel and a glass coverslip inducing sporozoites to move into the hydrogel. (B) Overlay of sporozoite tracks (black) and DIC image showing sporozoites inside a soft hydrogel at the end of a recorded image sequence. Scale bar, 20 μm. (C) Speed of WT sporozoites moving in the skin *in vivo* (37) and in hydrogels with small or large pores. (D, E) Speed of indicated mutant and control (Ctrl) parasites in hydrogels with small (D) and large (E) pores. (F, G) Speed of drug-treated and control parasites in hydrogels with small (F) and large (G) pores. Note the faster speed in gels with larger pores. Data from at least two independent experiments. Numbers above bars indicate the number of sporozoites analyzed. Significance determined by Mann-Whitney test.

### Diminished sporozoite motility on endothelial cells and soft substrates

Upon entry into the blood circulation of a vertebrate host, the sporozoite needs to travel to the liver without ‘loosing time’ or ‘getting lost’ by adhering to and migrating on the endothelial cells of the blood vessels or by crossing the endothelial cell layer and penetrating the surrounding tissue. How the sporozoites, which can migrate on virtually all biological surfaces, avoids doing so is unclear. One possibility might be the lack of a receptor on endothelial cells that specifically allows adhesion under flow. Alternatively, the relative softness of endothelial cells (27) might inhibit adhesion and/or motility. We therefore compared sporozoite motility on fibroblasts, endothelial cells and hepatocytes (**Figure 4 A, B**). Interestingly, sporozoites moved robustly on confluent layers of fibroblasts and hepatocytes whereas only a small fraction moved on monolayers of different endothelial cells (**Figure 4 C**). Strikingly, we also found that similar to endothelial cells sporozoites did not move on pericytes (**Figure 4 C**). This effect cannot be attributed to the different cell media, as sporozoites moved at similar high rates on glass in the different cell media. This suggests that sporozoites lack the capacity to migrate efficiently on cell forming blood vessels. To probe if this decrease of motility could indeed be due to substrate stiffness, we tested whether a physiologically relevant range of elasticities affects sporozoite motility. To this end we performed cell migration assays on soft, intermediate and stiff 2D hydrogels, with the soft and intermediate hydrogels reflecting the elasticity of endothelial cells and fibroblasts, respectively (**Figure 4 D, E**). Interestingly, the motile fraction of sporozoites significantly dropped from medium to soft hydrogels but remained constant between medium and stiff hydrogels (**Figure 4 F**). The number of circles traced by migrating sporozoites, a combined measure for persistent migration and speed, increased with increasing substrate stiffness (**Figure 4 G**). As substrate stiffness did not affect the speed of continuously moving sporozoites (**Figure 4 H**), the decreased stiffness diminishes persistence of movement. Together these data suggest that sporozoites are less capable to move for long periods of time on endothelial cells. This apparent lack could provide a key evolutionary advantage in allowing efficient transport of sporozoites to the liver and hence efficient propagation of the life cycle.

**Figure 4.**
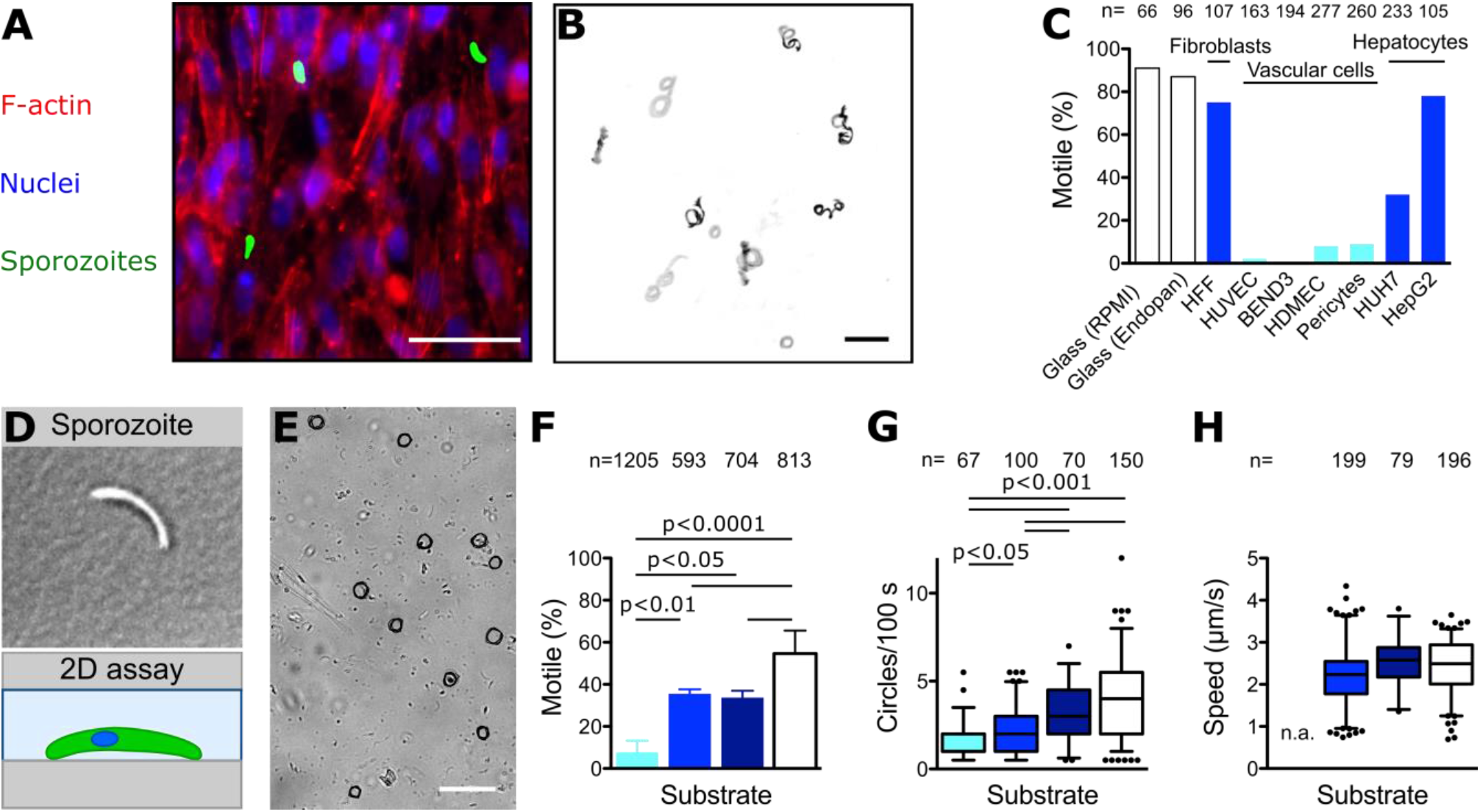
Sporozoite motility on cells and 2D hydrogels of different stiffnesses. (A) Merged fluorescence images showing sporozoites on cultured HFF cells. Actin filaments of HFF cells were stained with phalloidin (red), nuclei were stained with Hoechst (blue) and sporozoites expressed GFP (green). Scale bar, 50 μm. (B) Maximum projection of sporozoites moving on cultured HFF cells. Scale bar, 50 μm. (C) Percentage of motile sporozoites on glass (white bars) as well as confluent layers of fibroblasts, endothelial cells and hepatocytes. Soft endothelial cells indicated in cyan, others in blue. (D) Sporozoite on planar uncoated hydrogels. (E) Overlay of sporozoite tracks (black) and DIC image showing sporozoites on a hydrogel. Scale bar, 50 μm. (F) Percentage of motile sporozoites on hydrogels of different stiffnesses and glass 30 min after activation with BSA. Significance determined by One-way analysis of variance with Bonferroni’s Multiple Comparison test. (G) Migration persistance as determined by the number of circles performed within 100 s on hydrogels of different stiffnesses and glass 30 min after activation with BSA. All sporozoites moving for more than one parasite length within the time of observation were analyzed, even if they stopped moving or detached during imaging. Significance determined by Kruskal-Wallis test with Dunn’s Multiple Comparison test. (H) Speed of sporozoites moving consistently for at least 60 s on substrates of different stiffnesses. Note that not all sporozoites analyzed in (G) complied with that requirement. To be able to analyze a high number of sporozoites, movies were taken 10 to 45 min after activation with BSA. Speed of sporozoites on soft hydrogels. was not analyzed (n.a.) due to the low fraction of motile sporozoites on these hydrogels. No significant differences.

## DISCUSSION

### Ookinetes but not sporozoites display different modes of motility dependent on substrate elasticity

Here we show that ookinetes and sporozoites are able to move at fast speeds if confined between or within uncoated PA hydrogels. Tuning of these PA gels to elasticities as they occur on the natural substrates encountered by the parasites revealed intriguing differences between the motile behavior of ookinetes and sporozoites. While ookinetes moved at relatively slow speed for over 20 hours on soft gels, sporozoites moved at ten-fold higher speed for less than 1 hour. Furthermore, ookinetes adapted their migration path to substrate stiffness, while sporozoites either migrated on their typical circular paths or did not. Adaptations of migration paths in ookinetes might be an adaptation to the very different requirements the ookinetes have to master in their short life. Firstly, they need to exit the blood meal, a process that appears to depend on gravity in mosquitoes that agglutinate their blood meal, while it is independent of gravity in those that do not. The latter suggests, that active migration through the blood meal might well play a role in ookinete dissemination as diffusion is likely inefficient. Our results showing that ookinetes migrate as well on the softest possible gel as they do on the hardest substrate (glass), suggests that indeed their motility apparatus and shape might have evolved to allow active dissemination within the blood bolus. *In vivo*, this might not manifest itself as persistent migration but be limited to the occasional displacement of a red blood cell or the agglutinated debris thereof with the parasite progressing slowly across the lumen of the midgut towards the epithelial cells at the edge. Indeed active ookinete motility has recently been demonstrated within the blood meal by *in vivo* imaging (6). However, more long-term imaging needs to be performed to understand how this movement leads to ookinete displacement in the blood meal. The observation that ookinetes move in a predominately circular manner on the softest gels yet in a predominately linear fashion on harder surfaces suggests that the parasite is sensing the stiffness of its environment and adapts its motile behavior accordingly. This could be relevant as the parasite migrates from the soft blood meal through the less soft epithelium or as it seeks to arrest below the basal lamina, which is likely the least elastic substrate the ookinete encounters. To understand the physiological relevance of this switch in migration path, it would now be interesting to screen transgenic ookinetes with documented defects in migration for their capacity to undergo this transition from circular to linear motion and to correlate this with *in vivo* observations.

### A synthetic 3D environment to mimic sporozoite migration in the skin

We found that ookinetes do not move on planar uncoated PA hydrogels while sporozoites can move continuously on stiff PA hydrogels. This indicates that there is an important difference in parasite motility dependent on the dimensionality of the environment, which might be due to a different capacity to build adhesion sites or due to the arrangement of organelles. In contrast to ookinetes, sporozoites show a very clear chiral cellular architecture with strongly bend polar rings (4, 36). Our data implies that adhesive interactions are especially important for motility on planar substrates, while 3D motility of the parasites is irrespective of receptor-ligand interactions. Unlike ookinetes, sporozoites do get in contact with planar substrates (transition from hemolymph to salivary gland and blood circulation to liver) as well as 3D environments (skin) on their journey from mosquito to mammal.

Interestingly, we could find sporozoites moving within soft PA hydrogels, which feature narrow pores (30). Their speed inside these hydrogels was similar to the range of speeds observed *in vivo* in the skin (11, 12, 37) or natural hydrogels (32). Yet, their trajectories were more regular than in the skin, which might be explained by the heterogenous composition of the skin consisting of different extracellular matrix proteins, fibers and cell types. The robust migration within the gels shows that BSA is sufficient to activate 3D motility of sporozoites and, importantly, that there is no need for specific receptors. While such specific interactions between substrate proteins and sporozoite surface proteins clearly exist, they might only play minor modulating roles during migration as shown for TRAP (38). Indeed, leukocytes are able to move in the absence of integrins through confined environments, but not on planar substrates, suggesting that receptor-ligand interactions are less important for 3D motility of cells in general (39, 40).

To probe how well PA hydrogels can reconstitute sporozoite migration in the skin, we used two mutant parasite lines that show different *in vitro* and *in vivo* migration capacities. Interestingly, we found that confinement can fully compensate for the lack of the actin-binding protein coronin as was previously shown by *in vivo* imaging in the skin (33). We also observed similar speed ranges of wild type sporozoites and their susceptibility to motility inhibiting drugs. This suggests that the 3D PA hydrogels represent good platforms to investigate *Plasmodium* sporozoite migration *ex vivo* and might be useful for rapid drug and antibody testing.

### Sporozoites only move on stiffer substrates – a mechanism for avoiding premature adhesion?

Like ookinetes, sporozoites can also migrate on a range of different substrates. Motility on soft PA gels was first shown in a study that measured traction forces of migrating sporozoites (35) and raised the question how parasites link to their substrate. Several observations such as transient adhesions (35, 41), active processing of adhesive proteins (42) and sensitivity to lateral flow (21) suggest that sporozoites avoid strong attachment to their respective substrates during migration in order to achieve their high speed of more than 1 μm/s (12, 13, 43). However, sporozoites need to attach specifically to target organs such as the salivary gland or the liver. Clearly, a complex machinery must regulate this switch in behavior, likely involving calcium and cGMP signaling, adhesins as well as cytoskeletal arrangements (20, 41, 44–46). This switch from migration to adhesion and invasion does not have to be abrupt but could also be gradual. It is currently not known if sporozoites pass by the salivary glands several times and how frequently they attach to and detach from the endothelium in the liver sinusoids before they finally attach and enter the organs. Nano-patterning of ligands on non-adhesion permitting surfaces shows that only a few dozen adhesins are necessary to allow sporozoite motility (23, 47). This could suggest that the major surface protein CSP mediates low affinity unspecific binding to a surface with adhesins of the TRAP family providing for stronger but transient substrate adhesion (18, 35, 41, 42, 48). Clearly, modulation of adhesion and deadhesion cycles plays a complex role in motility and invasion. For example, *P. berghei* sporozoites lacking the actin binding protein coronin rarely attached to and glide on a flat substrate. This translates into a lower number of sporozoites entering the salivary gland. However, those that do and are injected into the skin move fine in this 3D environment and can invade the liver (33). This, and other evidence (49, 50) suggests that the salivary gland is a stronger barrier to sporozoite invasion than the liver.

But why did the parasites evolve such a curious way of migration and avoidance of stickiness? Two driving forces might play their part: first, a less-sticky surface might not be targeted by the complement system or antibodies. Second, the parasites avoid getting stuck before they reach their destination. Ookinetes could easily be trapped in the blood meal if they would attach too strongly to red cells and their digested remnants. Similarly, sporozoites are injected into the skin of a new host and enter the blood stream often at the periphery, e.g. at smelly, mosquito-attracting feet. While they need to migrate to cross the dermis and to enter the liver, they should not emigrate on the long route from the bite site to the liver sinusoids. A mix of hydrodynamic flow, absence of specific ligands and, as shown here, low adhesion to endothelial cells might combine to enhance the efficiency of homing to the liver (**Figure 5**).

**Figure 5.**
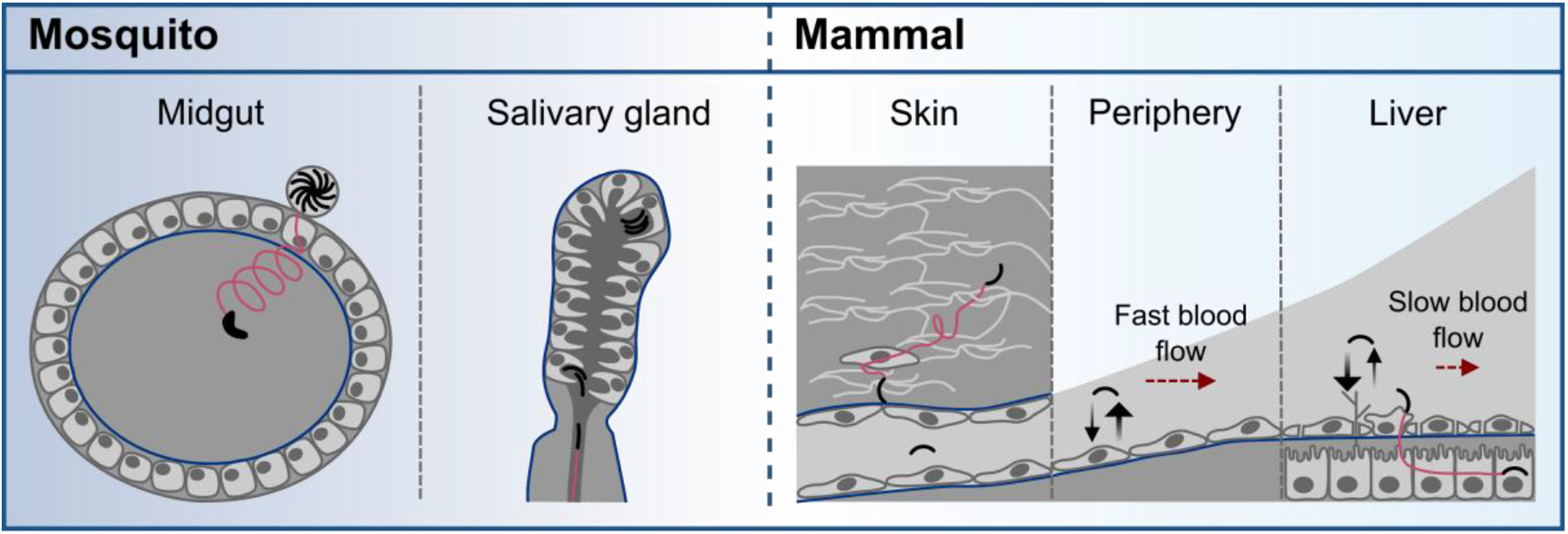
Scheme illustrating different modes of motility throughout the *Plasmodium* life cycle. Robust ookinete motility on soft substrates might allow for active dissemination within the blood bolus in the mosquito midgut with a change in migration mode upon contact of and passing through the epithelial cells. During an infectious bite, sporozoites need to pass through the narrow salivary duct to be transmitted into the mammalian skin. Once in the skin, sporozoites migrate and squeeze through pores to find and invade a blood vessel. With the blood flow (red arrow), sporozoites are transported through the body. The elasticity of endothelial cells does not favor adhesion (black arrows) so that sporozoites only arrest upon reaching the liver sinusoids where they escape the blood stream and infect hepatocytes (red line).

In conclusion, we show here that *Plasmodium* ookinetes migrate in different patterns on substrates of diverse stiffness, that sporozoites can migrate in PA hydrogels in ways mimicking their migration in the skin. We suggest that *Plasmodium* parasites evolved to finely tune their motility and adhesion machinery to avoid premature stickiness, which is important for efficient transmission to and from the mosquito vector of malaria.

## Materials and methods

### Ethics statement

Animal experiments were performed according to FELASA and GV-SOLAS guidelines and approved by the responsible German authorities (Regierungspräsidium Karlsruhe).

### Parasite culture, mosquito infection and parasite isolation

*P. berghei* parasite strains NK65 expressing GFP under the CS promoter, ANKA wt, *coronin(-)* and *hsp20(-)* were used (33, 34, 51). Ookinetes were cultured and purified as described before (50). Mosquitoes were infected and sporozoites isolated from salivary glands as described previously (8). Experiments with salivary gland sporozoites were performed 17 to 25 days post mosquito infection.

### Endothelial cell and pericyte cultures

Human umbilical vein endothelial cells (HUVECs; PromoCell), bEND3 and HDMEC were cultured in Endopan 3 Kit for endothelial cells (PAN-Biotech) supplemented with 10% FBS, 100 U/ml penicillin and 100 μg/ml streptomycin (both Gibco. by Life Technologies) in a 5% CO2 humidified incubator at 37C. Cells from passages 2 to 5 were used for the experiments. When indicated, cells were starved in growth factorfree Endopan 3 supplemented with 2% FBS, 100 U/ml penicillin and 100 μg/ml streptomycin.

Human pericytes (kindly gifted by P. Carmeliet) were cultured in alphaMEM glutamax, 100 U/ml penicillin and 100 μg/ml streptomycin, 10%FBS and 5ng/ml PDFGβ (R&D) and used until passage 7.

### Preparation of uncoated polyacrylamide hydrogels

PA hydrogels were prepared as described before (52). To tune the stiffness of the hydrogels (28), different monomer and crosslinker concentrations were added to the prepolymer solution. Soft gels were prepared with 5% AA/0.03% BIS, medium gels with 5% AA/0.3% BIS and stiff gels with 8% AA/0.48% BIS. To increase the pore size of soft gels, a gel formulation of 3% AA/0.06% BIS (small pores) or 3% AA/0.03% BIS (large pores) was used.

### Cell migration assays

A small volume of medium containing ookinetes was sandwiched between two hydrogels. After an incubation time of 5-10 min, medium was removed from the side to confine the ookinetes between the hydrogels. For extended imaging periods, the sandwich was sealed using paraffin wax. Alternatively, sporozoites were pipetted into a silicone chamber placed on top of a hydrogel. For 3D hydrogel assays, whole infected salivary glands were dissected into 30 μl of medium on a glass coverslip (22×22mm) placed on top of a microscope slide. Subsequently, the salivary glands were covered with a hydrogel. As control, ookinetes and sporozoites were imaged on glass as described previously (50). As sporozoite motility decreases with increasing time of incubation in BSA containing activation medium, but the sporozoites also need some time to settle on the substrate and we could not centrifuge them down when doing experiments with hydrogels, we imaged sporozoites after 30 min of incubation on the different substrates at low magnification. For imaging on cells, 5 x 10^4^ cells were seeded into 8-well Labtek chambered cover glass dishes. When cells reached at least 95% confluency the RPMI or Endopan medium was replaced by medium containing 3% BSA and 3 to 5 x 10^4^ freshly isolated sporozoites. Prior imaging dishes were centrifuged at 800 rpm for 5 min to allow sporozoites to adhere. Imaging was performed on an inverted Zeiss Axiovert 200 M microscope. Images of ookinetes were recorded every 20 s for 5-10 min, sporozoites were imaged every 3 s for 2-3 min.

### Analysis

Parasites were classified as motile if they moved for more than one parasite length during the time of observation. The percentage of motile sporozoites was determined by counting the total number of sporozoites within the first frame and dividing it by the number of parasites that moved at least half a circle as seen within the maximum projection of the image sequence. The speed was determined using the Manual Tracking plugin of Fiji (53). Only those parasites moving continuously for at least 60 s in the case of sporozoites or 5 min in the case of ookinetes were tracked. Mean square displacements of ookinetes were calculated from the position of the ookinetes in the first frame and in the following frames as given in the result files of the Manual Tracking plugin. Statistical analysis was performed in GraphPad Prism. Figures were generated using Inkscape.

## Acknowledgments

We thank Miriam Reinig for production of *Anopheles stephensi* mosquitoes, Rogerio Amino, Markus Ganter and Ulrich Schwarz for discussion and critically reading the manuscript. We would like to acknowledge the microscopy support from the Infectious Diseases Imaging Platform (IDIP) at the Center for Integrative Infectious Disease Research, Heidelberg, Germany. The work was funded by grants from the Human Frontier Science Program RGY0066/2015 (http://www.hfsp.org) to FF, by the FRONTIER program of Heidelberg University (http://www.uni-heidelberg.de/exzellenzinitiative/zukunftskonzept/frontier_de.html) to FF and CRA, the Deutsche Forschungsgemeinschaft (DFG, German Research Foundation) – Projektnummer 240245660 - SFB 1129 (http://www.sfb1129.de) and ANR-DFG grant FR2140/11-1 to FF. FF and CRA are members of the CellNetworks Cluster of Excellence at Heidelberg University. JR was member of the Heidelberg International Graduate School for the Biosciences (HBIGS) and XS was a member of the Master Program Molecular Biotechnology at Heidelberg University.

## Notes

### Competing Interest Statement

The authors have declared no competing interest.

