## Supplementary figures and images for "Malaria parasites differentially sense environmental elasticity during transmission"

### Supplementary Movie 1

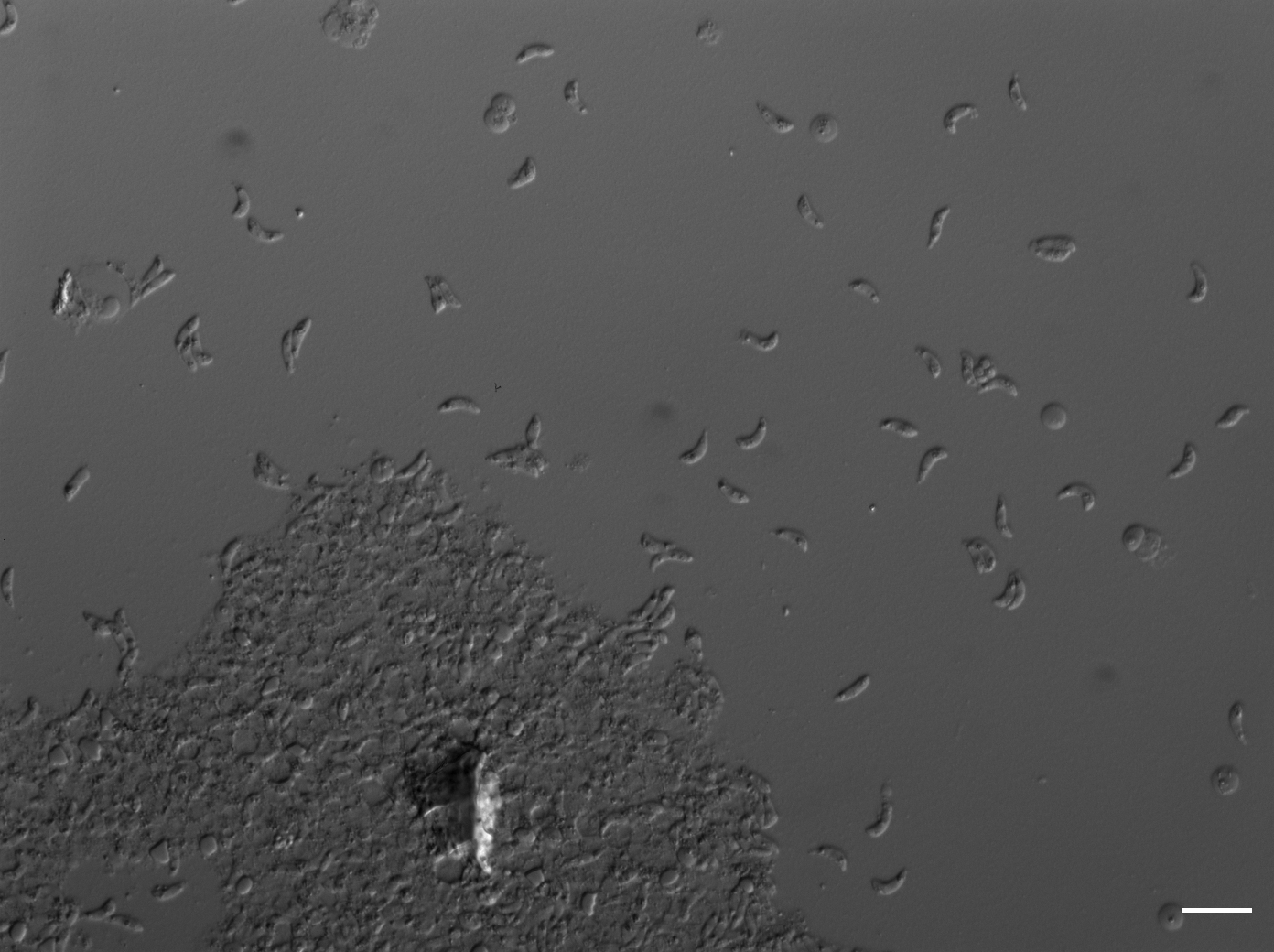

### Supplementary Movie 2

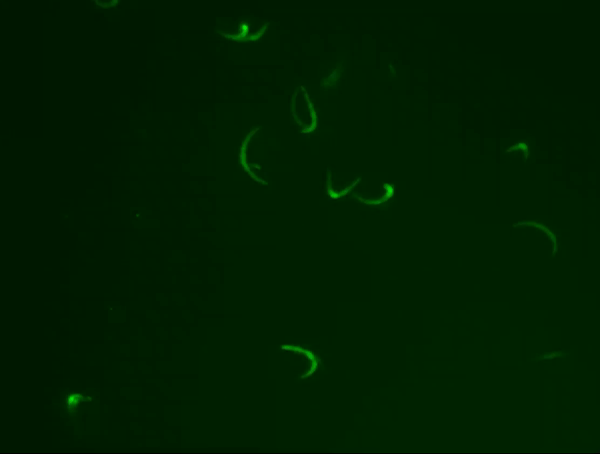
